# Efficacy of immunostimulant feed additive ShrimpGuard™, for the preventive management of White Spot Syndrome Virus (WSSV) infection in shrimps

**DOI:** 10.64898/2025.12.26.695863

**Authors:** E K Abhijith, Dhaval C Bamaniya, Suruchi Sharma, Indira Ghosh, Alagukanthasami Ponsrinivasan, Rahul Krishnan, Shanmukha Harish Kumar Paruvada, Saurav Kumar, Rishita Changede

## Abstract

White spot syndrome virus (WSSV) is a virulent lethal viral pathogen affecting shrimp aquaculture worldwide. Current solutions for WSSV management can be divided into three categories; farm management, selective breeding or genetic engineering to get WSSV resistant varieties of shrimp, and developing vaccine or drug-like solutions specifically targeting WSSV. At present, farm management or good farming practice is the most reliable way of managing the problem, anti-WSSV vaccine, or anti-WSSV drugs that are reliable and scalable in the market are not available. We have developed a feed additive named “ShrimpGuard” using our SOLAQ (Solutions for Agriculture and Aquaculture) platform that specifically provides Shrimp protection against the WSSV infection. Here we report protection observed in both laboratory and farm demonstrations, in *Penaeus vannamei*. Collectively, these studies show that ShrimpGuard does not cause any adverse effects on shrimp health and is able to provide significant protection to shrimp against WSSV infection. Briefly, the immune parameters of shrimp respond to ShrimpGuard treatment with increase observed in Total haemocyte count, and activity of prophenoloxidase. Further, survival rates of shrimp treated with ShrimpGuard after WSSV infection was between 80-100 % compared to 0% survival observed in untreated shrimp. The ShrimpGuard’s performance was tested in production farms and it was found that ShrimpGuard is safe for farm-based applications and the data obtained shows a positive impact on shrimp yield after ShrimpGuard treatment.

## Introduction

Shrimp aquaculture is one of the fastest growing protein production sectors due to its growing importance as a sustainable protein source. Global production of shrimp via aquaculture had reached 40 billion USD in 2019 (1). Among the different shrimp species used in aquaculture production, Pacific whiteleg shrimp (*Penaeus vannamei*) constitutes more than half of the global shrimp production (2). Intensification of shrimp farming has contributed to a greater prevalence of disease outbreaks, which in turn has driven the overuse of antimicrobial treatments resulting in a number of unintended consequences, including export rejects (3). Hence sustainable disease management solutions are required for disease management in shrimps. Among the various threats affecting the shrimp farming industry, viruses stand out as one of the most significant impediments to productivity where viruses account for 9 out of 10, among the causative agents of diseases that affect penaeid shrimp. They are subjects of study and are recognized as notifiable diseases by the World Organization for Animal Health (formerly OIE), which establishes international regulations for animal trade. In addition, viral infection can lead to secondary infections caused by bacteria such as Vibrio spp, leading to mortality or losses at the production farms. White spot disease (WSD), caused by white spot syndrome virus (WSSV), is one of the devastating viral disease affecting farmed shrimp worldwide (4, 5). WSSV causes mass mortalities up to 100% within 3-10 days of detection of the infection resulting in massive economic losses worldwide (6–9). WSSV has caused estimated economic losses of up to US$ 21 billion since its initial report in 1992 (10). For instance, a comprehensive economic loss study from India estimated annual loss of production due to WSSV to be 2.58 mt per hectare for a crop cycle with a corresponding loss US$ 240 million (11).

The WSSV is a large enveloped virus that has a double stranded DNA genome estimated to be 280 - 315 kb in length (12). It belongs to the family Nimaviridae & Genus Whispovirus (13, 14). Clinical symptoms of WSSV infection include rapid development of white spots (ranging from 0.5–3.0 mm in diameter) on the shrimp exoskeleton, appendages and inside the epidermis. Since these spots can also be produced by other factors like bacterial infection, stress, high alkalinity, they are not always reliable for preliminary diagnosis. Reliable diagnosis is performed by specific PCR based methods. Symptoms of WSSV infected shrimps include significant lethargy, sudden reduction in feed consumption and notable red discoloration of body and appendages (7). The problem is amplified as shrimps can eat other weak shrimps, thereby accelerating the spread of the virus in the pond (15). Hence, prevention and/or control of the disease in early stages is highly desired for the sustainability of the industry.

Due to its broad host and vector range, WSSV is efficiently transmitted within and between aquatic environments. Disease can spread through multiple vectors including, birds and other carriers (e.g. crabs, wild shrimps) that reside in the creek or ocean, which provides the water source used for water exchange in the farms. In addition to shrimps, WSSV has a wide range of hosts or carriers with over 93 species of arthropods such as such as salt, brackish and freshwater crayfishes, crabs, lobsters (16). The virus is highly transmissible and infects all cultured penaeid shrimps. WSSV can be transmitted to healthy, susceptible shrimp, within or between shrimp farms, by (i) dead or moribund shrimp, (ii) via the contaminated water sources or (iii) from infected shrimp, brooders or post-larvae (17). Vertical transmission takes place when viral particles shed during spawning are ingested by larvae at first feeding; however, it remains uncertain whether WSSV virions are contained within the shrimp eggs themselves (14). Horizontal transmission of WSSV can also occur through sediments & carrier organisms. Because of its broad host range, WSSV is not only a major threat to shrimp farming but also to the worldwide marine ecology (18). Hence, immediate measures to control it are essential.

Broadly the current solutions for WSSV management can be divided into three categories; farm management, selective breeding to get WSSV resistant shrimp varieties, and developing anti-WSSV vaccine or anti-WSSV drug like solution specifically targeting WSSV. At present the farm management or good farming practice is the most reliable way of managing the problem. Different strategies have been tried and tested to boost the shrimp’s immunity against harmful pathogens during farming like introduction of natural immunostimulants including probiotics, phytobiotics, organic acids etc. as feed additives (19). Various studies have shown that shrimps do not possess a canonical adaptive immune response as observed in vertebrates. However, there is evidence that suggests a prior exposure to the antigen of a pathogen provides protection from subsequent infection by the pathogen even in invertebrates. This protection is referred to as specific-immune priming. Precise mechanisms for specific-immune priming in invertebrates are not fully elucidated; however, there are reports that provide insights about the possible mechanisms for this type of response (20–23). Relying on the presence of a high degree of specific-immune priming in shrimps, various studies have been conducted to vaccinate shrimp via injection or oral delivery of pathogen proteins (24–29). The vaccination has been attempted using various approaches that include sub-unit protein vaccines, whole virus inactivated vaccines, DNA vaccines, RNA vaccines and RNA interference approaches which has demonstrated protection against the pathogen (30).

Even though different approaches for managing WSSV have been tried and tested, there is still a need to identify a scalable product/mechanism for managing WSSV in field conditions. Multiple solutions can be used together to combat the threat of this virus. In this study, we present a feed additive named “ShrimpGuard” that has been developed using our SOLAQ platform.

SOLAQ is our cutting-edge proprietary platform that delivers targeted disease management solutions for aquaculture. Our platform integrates three core disciplines: bioinformatics, synthetic biology & material science to design and deliver safe, orally deliverable prophylactic and therapeutic products. Our proprietary bioinformatics engine enables precise identification of pathogen-specific targets. These insights fuel our synthetic biology workflow to generate customized biologics for a specific disease threat. Finally, advanced material science ensures these biologics are effectively encapsulated for oral delivery, ensuring stability, ease of storage and practical on-farm use.

ShrimpGuard contains novel peptide-based biologics that specifically activate the shrimp innate immune system against WSSV, provides protection against infection, extends culture time, and reduces the risk of mortality due to WSSV. This study reports the efficacy of ShrimpGuard in protecting shrimp against WSSV observed in laboratory and field conditions. Collectively, these trials show that ShrimpGuard boosts immunity, does not cause any adverse effects on shrimp health, and is able to provide protection to shrimp against WSSV infection.

## Materials & Methods

### Shrimps for laboratory trials

Specific pathogen free (SPF) shrimp (P. vannamei) utilized in this trial were obtained from the Shrimp Improvement Systems in Hawaii, U.S.A (SIS Hawaii) broodstock (31–33). Post Larvae (PLs) were produced under strict bio-security in designated hatcheries and screened for all important pathogens, including White Spot Syndrome Virus (WSSV), Infectious Hypodermal and Hematopoietic Necrosis Virus (IHHNV), Acute Hepatopancreatic Necrosis Disease (AHPND) (34) using PCR and real-time PCR prior to shipping to Laboratory for safety and efficacy trials. At the Laboratory, the PLs were reared in primary quarantine tanks for 7 days. During the primary quarantine, the PLs were checked for the above-listed disease using PCR to confirm the disease-free status of the stock and absence of *Enterocytozoon hepatopenaei* (EHP) (35). The PLs which passed the primary quarantine were cultured in strict bio-secure conditions for secondary quarantine for 30 days. The juveniles were checked again for the listed diseases using PCR techniques.

### Shrimp feed and maintenance

Shrimp were fed with a standard commercial feed diet (without any additives) at satiation for 4 meals per day during the trial (5% estimated body weight). The amount of feed was adjusted depending on the estimated biomass and shrimp feeding behaviour of the tanks.

### Water quality measurement and maintenance

Water quality parameters such as temperature, dissolved oxygen (D.O.), pH, salinity, total ammonia nitrogen (TAN), nitrite, alkalinity, K^+^, Ca^2+^, Mg^2+^ were maintained at optimal ranges for shrimp normal growth (33). After the tests, water was chlorinated for 24 hours and then discarded.

### WSSV inoculum for infection

#### Injection

For injection efficacy trials, the shrimps were challenged by WSSV per os challenge method (36–38). For preparation of the inoculum, WSSV-clarified extract was mixed with a commercial diet (without prophylactic treatment) at the ratio of 1:3 (weight/weight) and kept at −80°C until use. The inoculum had an average WSSV load of 0.5-1.0×10^7^ copies/g inoculum and expected to kill 50% shrimp in 156-173 hours (Lethal dose 50, 156-173 hours, which had been determined prior to the start of the trial and further confirmed using probit analysis (39) in the Positive control samples).

#### Oral

For oral efficacy trials, the fresh WSSV inoculum was prepared from WSSV-infected shrimp by performing WSSV infection on live shrimp using the intramuscular injection method. The muscle of infected shrimp was homogenized in Phosphate Buffer Saline (PBS) at the ratio of 1:9 (40). The homogenate was centrifuged at 6000 g for 10 minutes at 4°C and the supernatant was filtered through a 0.45-micron filter. The filtered supernatant was kept in the tube and stored in −80°C freezer.

#### Oral trial - injection challenge

The shrimp in both the positive control and treatment groups were challenged with purified WSSV inoculum at a dose of 1 x 10^4^ copies/50 μL per shrimp, administered via intramuscular injection. The intramuscular challenge ensured uniform viral delivery compared to the per os challenge (oral route of infection), allowing for effective evaluation of treatment efficacy as described in previous studies (36).

#### Oral trial - per os challenge

The shrimp in the negative control were fed non-WSSV feed, and shrimp in the positive control and treatment groups were fed the WSSV-coated feed using the per os method described by Lightner, D.V. et al (36–38, 41). In this trial, WSSV-coated feed was fed to the shrimps in the challenged groups 3 meals per day and basal feed was fed as the remaining one meal per day. Mode of transmission of WSSV is oro-fecal route and this method allows for infection to proceed naturally, though the pathogen load per animal cannot be controlled. This allows is to assay whether protection would be effective when the virus enters through the oral route.

### Experimental Design

#### Safety trial

A completely randomized trial (CRT) was designed using 90 L tanks. Shrimps were divided into four treatment groups: Group-1, Group-2, Group-3, and Group-4. Each treatment group comprised three tanks; with n shrimps in each tank (n varied among trials and is stated in the results). The weight of individual shrimps in each tank was recorded as the initial shrimp weight. Group-1 served as the negative control, while Groups 2, 3, and 4 were used to test TW-1, TW-2, and TW-3 biologics, respectively.

#### Efficacy trial

A completely randomized trial (CRT) was conducted using experimental tanks. Shrimps were divided into five treatment groups: Group-1, Group-2, Group-3, Group-4, and Group-5. Each group consisted of three replicate tanks, with 6 to 20 shrimps per tank (number varied among trials, as trials were conducted at different laboratories and is described in the results). Group-1 served as the negative control, Group-2 served as the positive control, while Groups 3, 4, and 5 were used to test the TW-1, TW-2, and TW-3 biologics, respectively.

#### Administration of TW Biologics

For the injection-based trials (safety and efficacy), the biologic formulation was administered via intramuscular injection at a dosage of 6 μg per gram, 6 μg of TW biologics per gram of the total formulation as per respective trial plan. Control groups received an equivalent volume of sterile phosphate-buffered saline (PBS). All animals were monitored post-administration for signs of stress, abnormal behavior, or mortality to evaluate injection safety prior to challenge or performance assessment.

For the oral trials (safety and efficacy), the biologic was incorporated into the feed at a final concentration of 1% TW biologics per feed, 10 g of TW biologics in 1 kg of feed. The supplemented feed was air-dried and stored at room temperature until use. Feeding was conducted at a rate of 5% of body weight per day as per respective trial plan. Control groups received the same basal feed without biologics. Shrimp were observed daily to assess feed acceptance, general health, and survival throughout the trial.

The experimental duration and sampling framework for each respective trial are outlined in the Results section.

### Expression analysis of immune related genes by qPCR

The expression of four immune-related genes - Prophenoloxidase, Toll-like receptor, C-type lectin, and Crustin—was quantified in experimental shrimp using quantitative realtime PCR (qRT-PCR). These genes were selected because they represent key components of the shrimp’s innate immune system, encompassing various immune response mechanisms, including pathogen recognition (Toll-like receptor, C-type lectin), immune signaling (Toll-like receptor), and effector defense mechanisms (Prophenoloxidase, Crustin) (29, 42–44).

Shrimp tissue samples from gills were dissected and preserved in TRIzol™ reagent (Thermo Fisher Scientific, USA) and stored at 80 °C were used for total RNA extraction following the manufacturer’s protocol. The extracted RNA was then reverse transcribed into complementary DNA (cDNA) using random hexamer primers. The resulting cDNA served as the template for quantitative PCR (qPCR) amplification, employing primer sets specific to the genes of interest. The gene-specific primer sequences and expected product size of all genes are listed in Table 1. After amplification, melting curve analysis was performed for all samples to verify the specificity of the amplified product. The correct product was confirmed by both its characteristic melting temperature and band size on the gel.

**Table 1.**
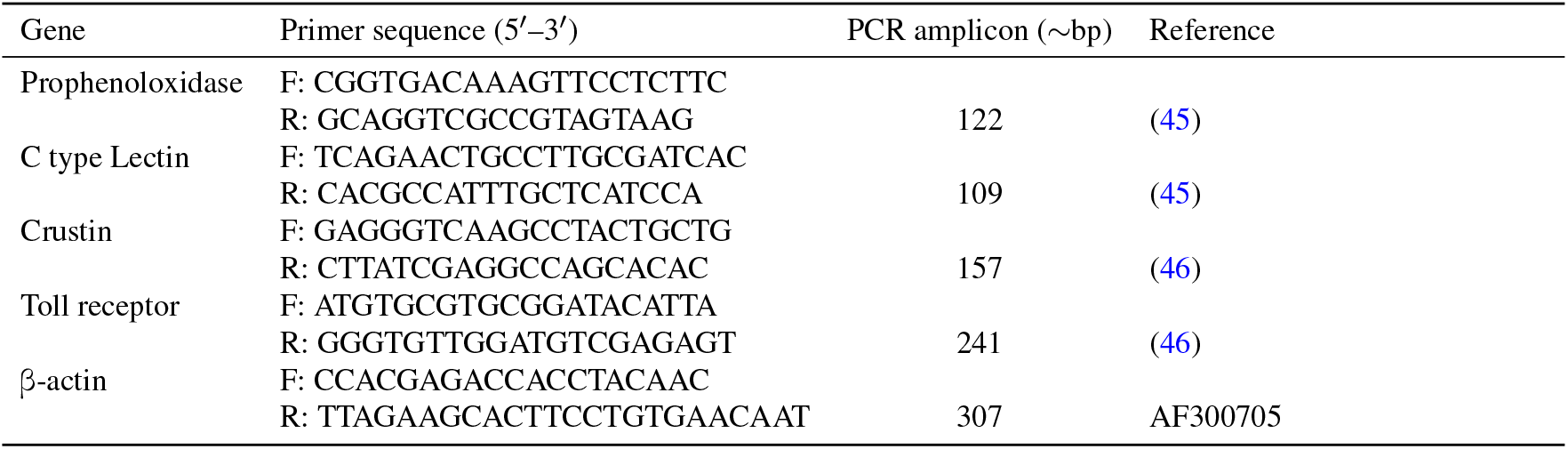
Primer sequences of the immune-related genes selected for the study.

All qPCR reactions were performed in technical replicates, and the mean cycle threshold (Ct) value from these replicates was used for subsequent analysis. The relative expression of each gene was calculated using the comparative Ct method (47), expressed as:

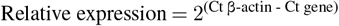

where β-actin served as the internal control. For each gene, the relative expression values from shrimp samples collected from the same pond were pooled, and the average relative expression value for each pool was calculated.

### Field trials and estimation of WSSV Copy Number

Samples were collected from each treatment pond at designated time points during the field trial to evaluate WSSV viral load dynamics. From each pond, pleopod and gill tissues were aseptically dissected from 10 representative shrimp using sterile instruments. The tissues were pooled and divided equally into two tubes to serve as duplicate pooled samples, immediately preserved in 95% molecular-grade ethanol (with ethanol replaced once after 24 hour), and stored at –20°C until DNA extraction.

Genomic DNA was extracted using the commercial extraction kit (Qiagen, Germany) following the manufacturer’s protocol. Quantification of WSSV copy number was performed by quantitative real-time PCR (qPCR) using the WSSV-specific primers RT-WSSV-F154 (5-CCAGTTCAGAATCGGACGTT-3) and RT-WSSV-R154 (5-AAAGACGCCTACCCTGTTGA-3) and a TaqMan probe labeled with FAM (5) and DABCYL (3) as described by Jang et al. (2009) (48). The viral copy number in each sample was determined by comparing Ct values to a standard curve generated from 10-fold serial dilutions of plasmid DNA containing the target fragment as described in Phuthaworn et al., 2016 (49). Viral copy numbers presented represent the mean of duplicate pooled samples per pond.

### Oral Delivery Demonstration – Evaluation of Biomaterial Absorption

A three-day feeding trial was conducted in *Penaeus vannamei* to assess the delivery efficiency of the Teora biomaterial. For this experiment, Teora biomaterial containing red fluorescent protein (RFP) was used to enable visualization of intestinal uptake. Two tanks were used for the study: one control tank receiving uncoated commercial feed and one treatment tank receiving feed coated with the RFP-tagged Teora biomaterial. Each tank was stocked with healthy shrimps of similar size and maintained under identical husbandry conditions throughout the trial. Shrimp in both groups were fed their respective diets with 5% of the body weight for three days, and no mortality was observed during the experiment. At the end of the trial, gut tissues were dissected, fixed with 10% formalin and examined under a fluorescent microscope to evaluate biomaterial uptake. Fluorescence intensity and distribution in the intestinal lining were visually assessed to determine the efficiency of biomaterial absorption in treatment compared to the control.

### Statistical Analysis

The survival rate and mortality rates of shrimp (%) were transformed using the arcsine-square-root transformation prior to analysis. Transformed data were analyzed by one-way analysis of variance (ANOVA), followed by Tukey’s multiple comparison test (if ANOVA indicated significance) to determine differences among treatments (50). All tests were two-tailed and significance was accepted at p < 0.05. Results are presented as mean ± standard deviation (51). All statistical analyses were performed using GraphPad Prism version 10.6.1.

## Results

### Injection Trials

Using our SOLAQ platform, we designed three active biologics and formulated them to provide shrimp protection against WSSV infection. To select the best-performing candidate from those designed by our platform for shrimp protection against WSSV infection, the active ingredients were synthesized and directly injected into the shrimp to evaluate their efficacy and safety.

#### Injection Trial: Safety Trials

First, the safety of the biologics was assayed. In this trial, each tank contained 20 shrimps per replicate, 3 replicates per treatment group. For the same, the shrimp were acclimatized in the trial tanks for two days. After acclimatization, Group-1 shrimps were injected with a plain buffer that did not contain any active ingredients, while Groups 2, 3, and 4 were injected with different TW biologics. After injection, the shrimps were maintained on a normal diet and observed for 7 days for any gross morphological defects or mortality. A schematic of the trial plan is shown in Fig. 1A.

**Figure 1.**
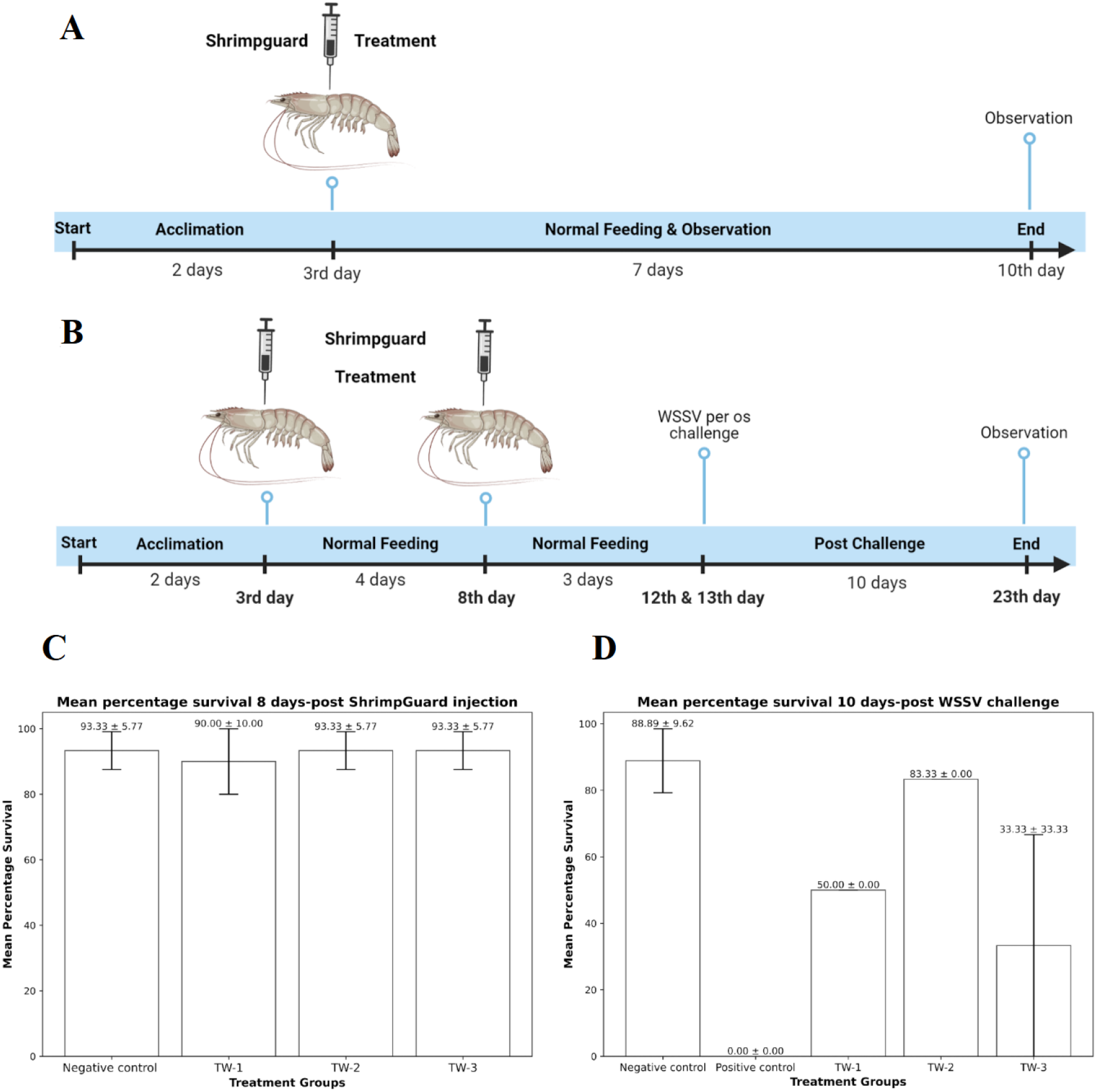
Safety and efficacy assessment of TW biologics administered via injection. A) Trial plan for safety trial. B) Trial plan for efficacy trial. C) Percentage survival of shrimps after 10 days of injecting with TW biologics. D) Percentage survival of shrimps 10 days after infection with WSSV in the indicated treatment groups. Negative control group was not infected with WSSV. The error bars represent the standard deviation of % survival in three independent different tanks of a given shrimp group.

The survival rate of Group-1, which was not injected with TW biologics, was 93.33 ± 5.77%. The survival rates for Groups 2, 3, and 4 were 93.33 ± 5.77%, 90.00 ± 0.00%, and 93.33 ± 5.77%, respectively (p > 0.05), as shown in Fig. 1C. The results indicate no statistical significance between the negative control and the treatment compounds, suggesting that the designed biologics, even when directly injected, are safe for shrimp.

In addition to assessing the safety of the TW biologics by survival rate in shrimp, the immune response was also evaluated to determine the treatment’s broader impact on shrimp health. Immune parameters, such as the Total Hemocyte Count (THC) & Prophenoloxidase (ProPO) activity were monitored at different time points 3, 7 & 14 dpi to assess whether the treatment elicited an appropriate immune response without adverse effects, as shown in Fig. 2A & Fig. 2B. Both THC and ProPO activity showed an increase in groups treated with TW biologics compared to the control group. This dual approach ensures that the treatment is safe when administered via injection and also supports an optimal immune profile, which is essential for the shrimp’s resilience against pathogens.

**Figure 2.**
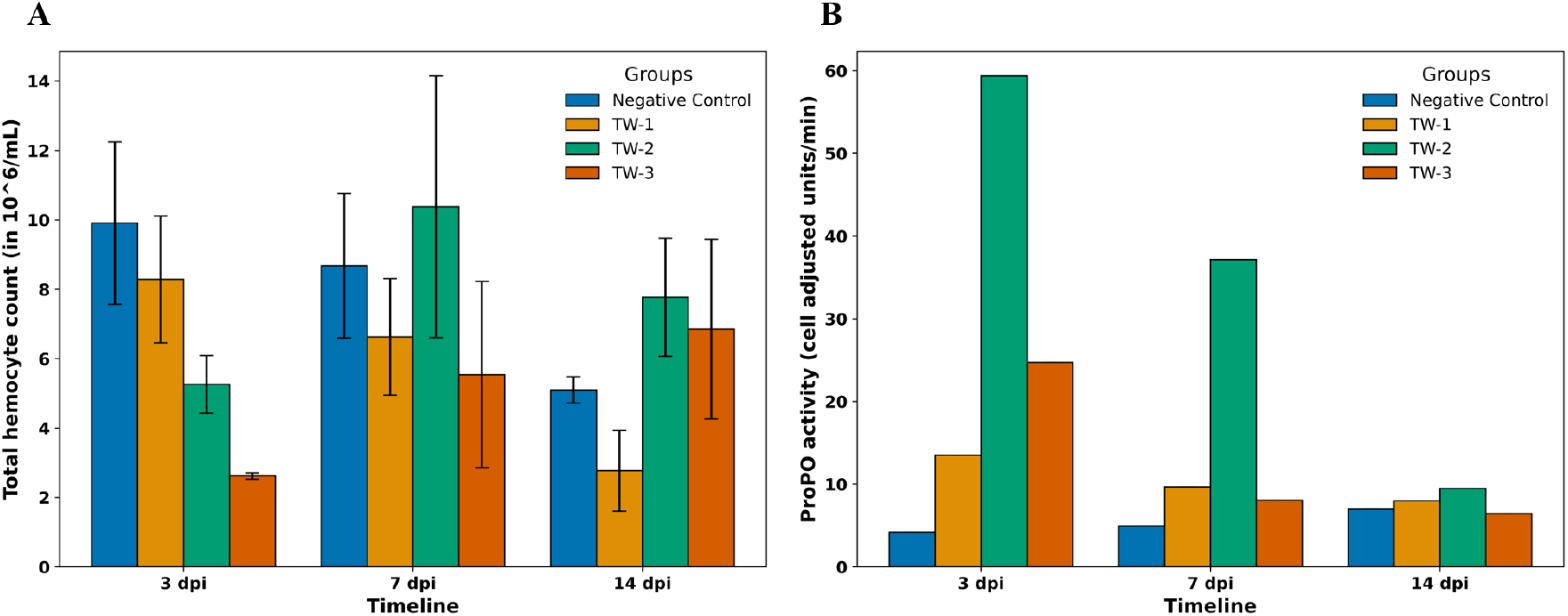
Immune parameters monitored from safety assessment of TW biologics administered via injection. A) Total hemocyte count (in 10^6^/mL) of shrimp treated with TW biologics monitored at 3, 7, & 14 dpi. B) Intrahemocytic ProPO activity (cell adjusted units/min) in shrimp treated with TW biologics monitored at 3, 7 & 14 dpi.

#### Injection Trial: Efficacy Trials

Next, we tested the efficacy of the biologics in a challenge trial. In this trial, each tank contained 6 shrimps per replicate. The shrimps were acclimatized in trial tanks for 2 days. After acclimatization, on day 3, Group-1 & Group-2 shrimps were injected with a plain buffer without any biologic, Group-3, Group-4, and Group-5 were injected with respective TW biologic. After the primary dose, shrimps were maintained under a normal diet for 4 days. On the 8th day of the trial, a booster or second dose of the respective biologic was injected in each group. After the secondary or booster dose, shrimps were monitored and maintained for 3 days under a normal diet. On the 11th & 12th day of the trial, all groups except Group-1 were infected with WSSV by per os challenge method (36–38, 41). The schematic of the trial plan is shown in Fig. 1B. The survival of shrimps was monitored for 10 days post-challenge. After 10 days postchallenge, the survival rate of Group-1, which was not infected with WSSV, was 88.89 ± 9.62%. Group-2, which was infected with WSSV but not injected with TW biologics, experienced 0% survival or 100% mortality. The survival rates of Groups 3, 4 & 5 were 50.00 ± 0.00 %, 83.33 ± 0.00 % & 33.33 ± 33.33 % respectively, as shown in Fig. 1D. Statistical analysis revealed a significant difference among the groups (one-way ANOVA, p = 0.0005). Tukey’s multiple comparison test showed that the survival rate of shrimps in TW-1 (p = 0.0175) and TW-2 (p = 0.0012) groups was significantly higher compared to the Positive Control group, whereas no significant difference was observed between TW-3 and the Positive Control (p = 0.134) after 10 days post-challenge under lab conditions. Among the TW biologics, shrimps treated with TW-2 biologic showed the highest survival rate with 83%. Based on this result we conclude that TW-2 is the best performing solution. However to validate this further, all TW biologics were converted into formulations that can be coated on feed and administered orally. These were then tested for efficacy and safety in subsequent oral trials. These trials demonstrate three things i) the best performing biologic is TW-2 (ii) not all biologics perform equally. This indicates that there is specificity in the biologic and this assay is able to select the best performing biologic, and (iii) this is not a nonspecific effect of encapsulation methodologies, which will be tested again in through oral delivery of the biologics. Hence, we next tested all the biologics by delivering them orally.

### Oral Trials

Our goal was to develop a product for shrimp administered orally through feed to protect against WSSV. We converted the best active ingredients into our proprietary oral formulation, that can be can be top coated onto normal feed using suitable binders, similar to nutritional supplements added at the farm. First, we demonstrated that our proprietary oral formulation can effectively deliver biologics in shrimp system. We then evaluated the safety and efficacy of TW biologics in laboratory and field trials.

#### Oral Delivery Demonstration

To evaluate whether orally delivered Teora biomaterial could be successfully absorbed through shrimp intestinal epithelium, gut samples from control & treatment groups were examined under fluorescence. The fluorescent microscopy analysis revealed clear differences in biomaterial absorption between the control and treatment groups. In the guts of shrimp from the control group, no biomaterials were absorbed. This can be seen under the red fluorescence image Fig. 3A, where there was no clear red fluorescence seen gathering at the intestinal lining. In contrast, shrimp from the treatment group showed evident absorption of the biomaterial. Under the red fluorescence image Fig. 3B, the intestinal epithelial lining can be clearly seen and defined by red fluorescence, suggesting that the biomaterial that was coated in the feed that contained a special red fluorescence protein to it was absorbed by the intestinal lining.

**Figure 3.**
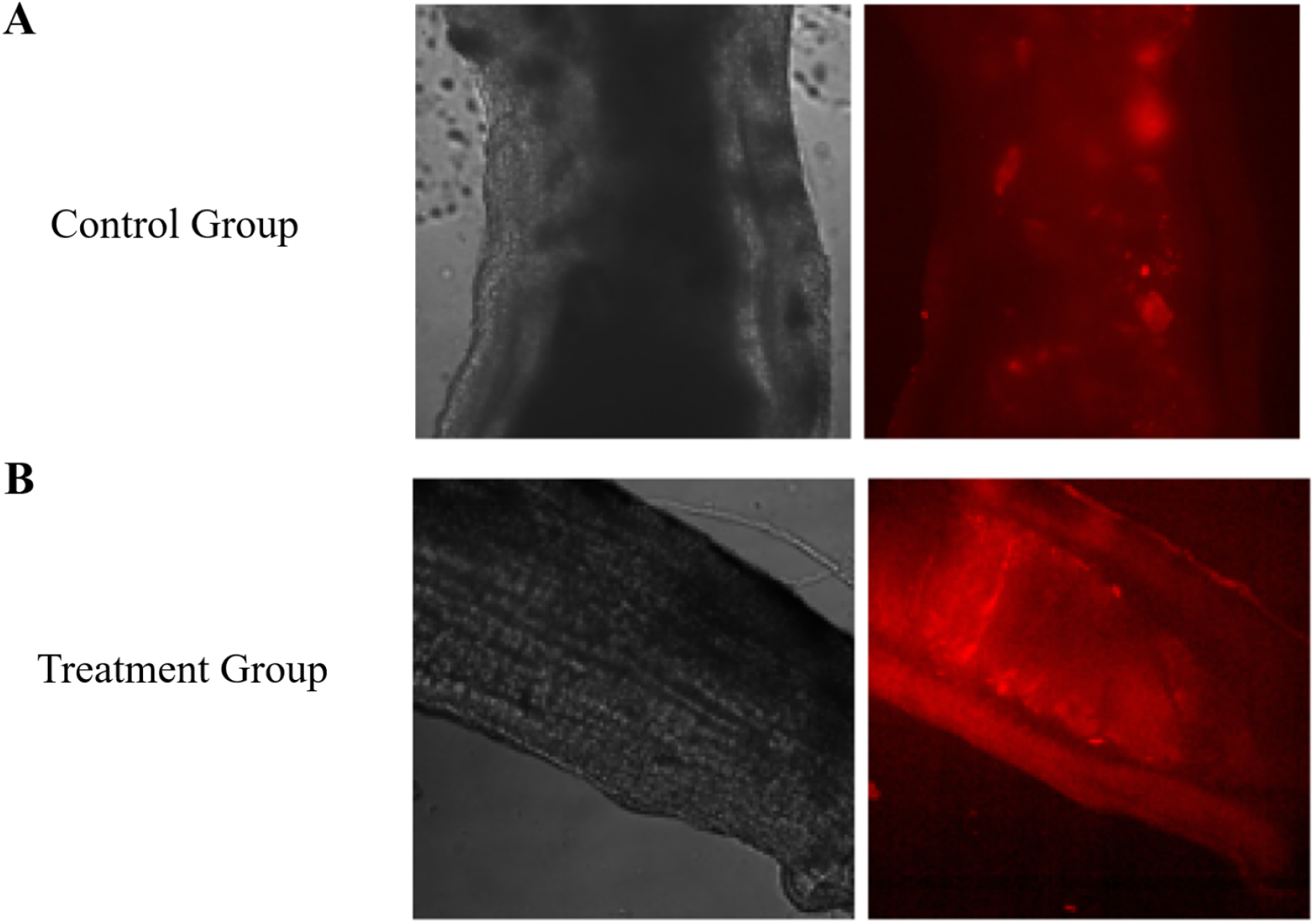
Bright field & Red fluorescence imaging of intestinal uptake of orally delivered Teora biomaterial in *Penaeus vannamei*. The left image shows a bright field view of shrimp gut under white light and right image shows the same gut region viewed under red fluorescence. A) Control group – Left image shows normal intestinal morphology with no visible biomaterial. Right image shows the same region where no detectable fluorescence signal is present, indicating absence of RFP tagged Teora biomaterial. B) Treatment group – Left image shows normal intestinal morphology with no visible biomaterial. Right image shows the same region, where the intestinal epithelium is clearly outlined by red fluorescence, confirming successful absorption of Teora biomaterial delivered coated via feed.

#### Oral trials: Safety Trials

In this trial, each tank contained 6 shrimps per replicate. The shrimps were acclimatized in tanks for 1 day. After acclimatization, Group-1 shrimps were fed with normal feed, while Groups 2, 3 & 4 were fed with feed top-coated with 2% TW biologics - TW-1, TW-2 & TW-3, respectively, for 5 days. Throughout the trial, shrimps were fed respective diets for 4 meals per day (approx. 5–10% of the estimated shrimp biomass). All the groups were monitored for 8 days for any morphological changes or mortality. A schematic of the trial plan is shown in Fig. 4A. After 8 days of safety trial, there was zero mortality across all the treatment groups as shown in Fig. 4B. These results indicate that the designed TW biologics delivered through our proprietary oral formulation are safe for shrimp. In addition, no reduction was observed in feed conversion ratio (FCR) between the treatment and control groups, suggesting that the formulation did not affect feed utilization efficiency.

**Figure 4.**
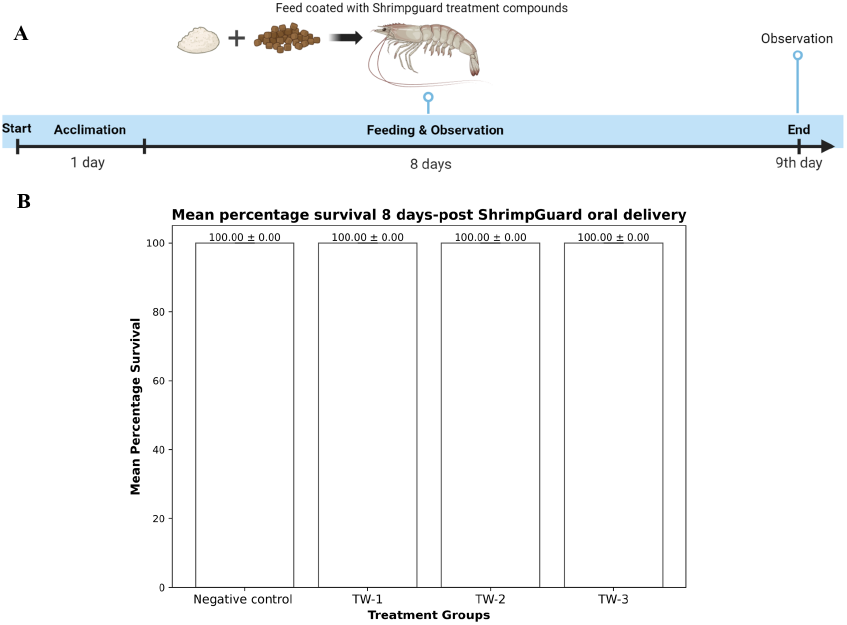
Safety assessment of ShrimpGuard™ administered orally. A) Trial plan for oral safety trial. B) Percentage survival of shrimps after 8 days of treatment with TW biologics administered orally.

#### Oral trials: Efficacy trials with intramuscular challenge

In this trial, each tank contained 15 shrimps per replicate. After acclimatization, Group-1 & Group-2 shrimps were fed with normal feed, while Groups-3, 4 & 5 were fed with feed top-coated with 2% TW biologics - TW-1, TW-2 & TW-3 respectively for 10 days. On the 11th day of the trial, all groups except Group-1 were infected with WSSV by intramuscular injection challenge (36). After the WSSV challenge, all groups were monitored and maintained for 20 days under normal diet to evaluate survival. The schematic of the trial plan is shown in Fig. 5A. After 20 days post-challenge, the survival rate of Group-1, negative control, which was not infected with WSSV and not fed with TW biologics, was 100%. The survival rate of Group-2, which was infected with WSSV but not fed with TW biologics, positive control, was 18.94 ± 2.58.

**Figure 5.**
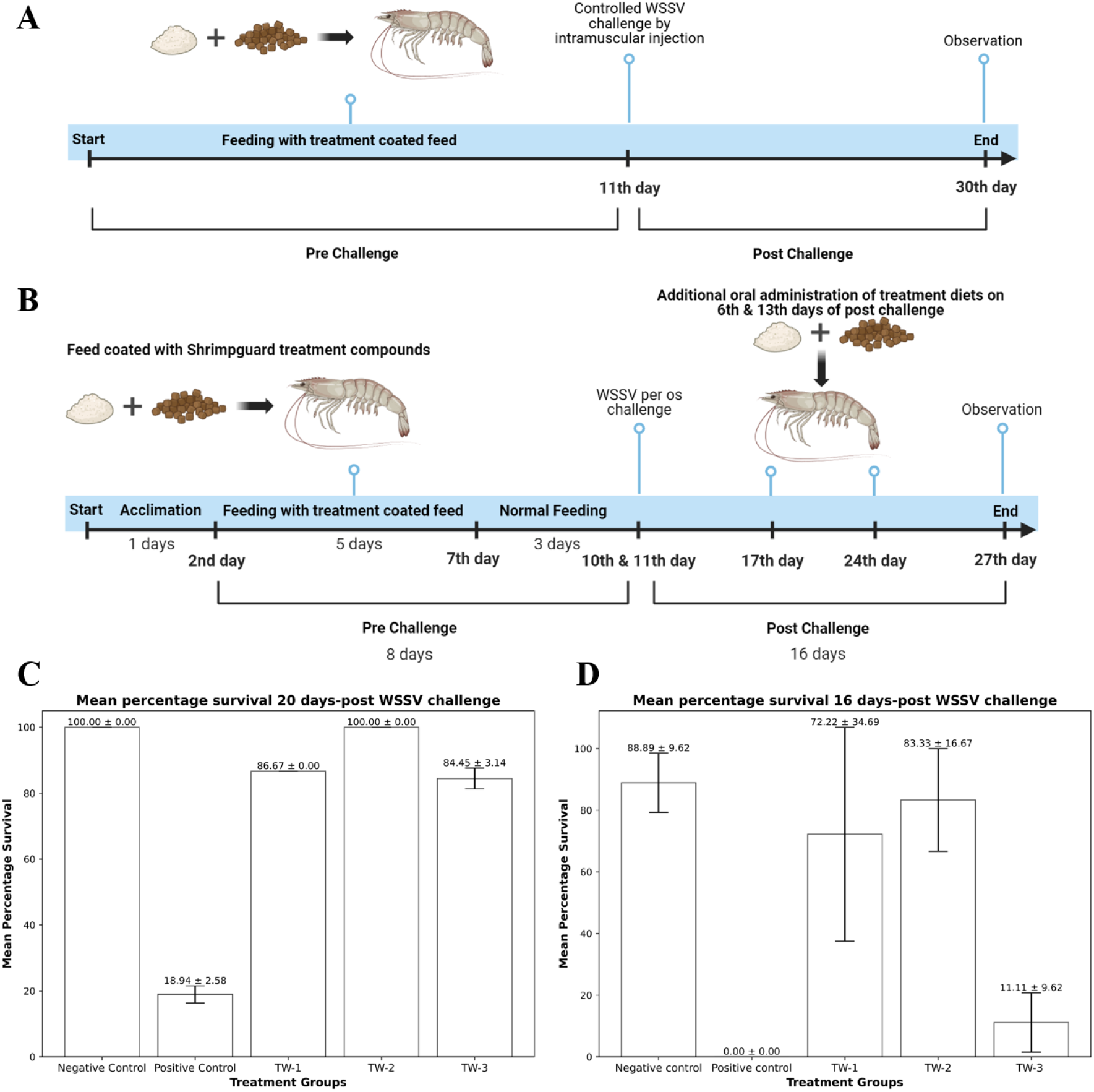
Efficacy assessment of ShrimpGuard™ administered orally against WSSV using different challenge methods. A) Trial plan oral efficacy trial with controlled WSSV challenge. B) Trial plan oral efficacy trial with per os WSSV challenge. C) Percentage survival of shrimps 20 days after infection with WSSV by intramuscular injection in the indicated treatment groups. D) Percentage survival of shrimps 16 days post WSSV infection in the indicated treatment group. Negative control group was not infected with WSSV. The error bars represent the standard deviation of % survival in three independent different tanks of a given shrimp group.

**Figure 6.**
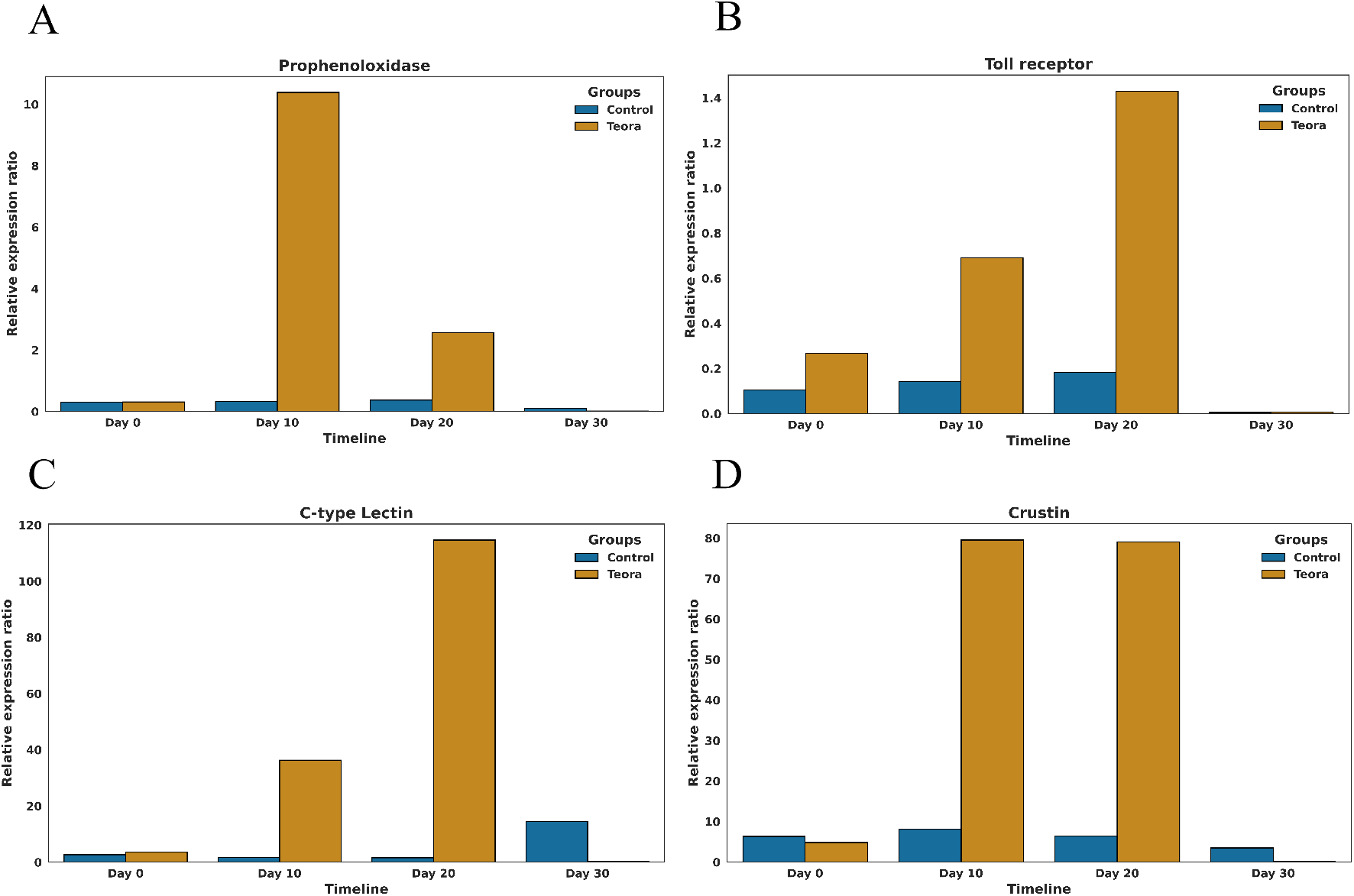
RT-qPCR analysis on the expression levels of immune genes in shrimp from the field trial. Relative expression ratio of A) Prophenoloxidase, B) Toll-like receptor, C) C-type lectin & D) Crustin. The gene expression levels were normalized with beta-actin. The bar graphs represent the relative expression ratio of shrimp from field trial ponds treated with ShrimpGuard™ compared to control ponds not treated with ShrimpGuard™

**Figure 7.**
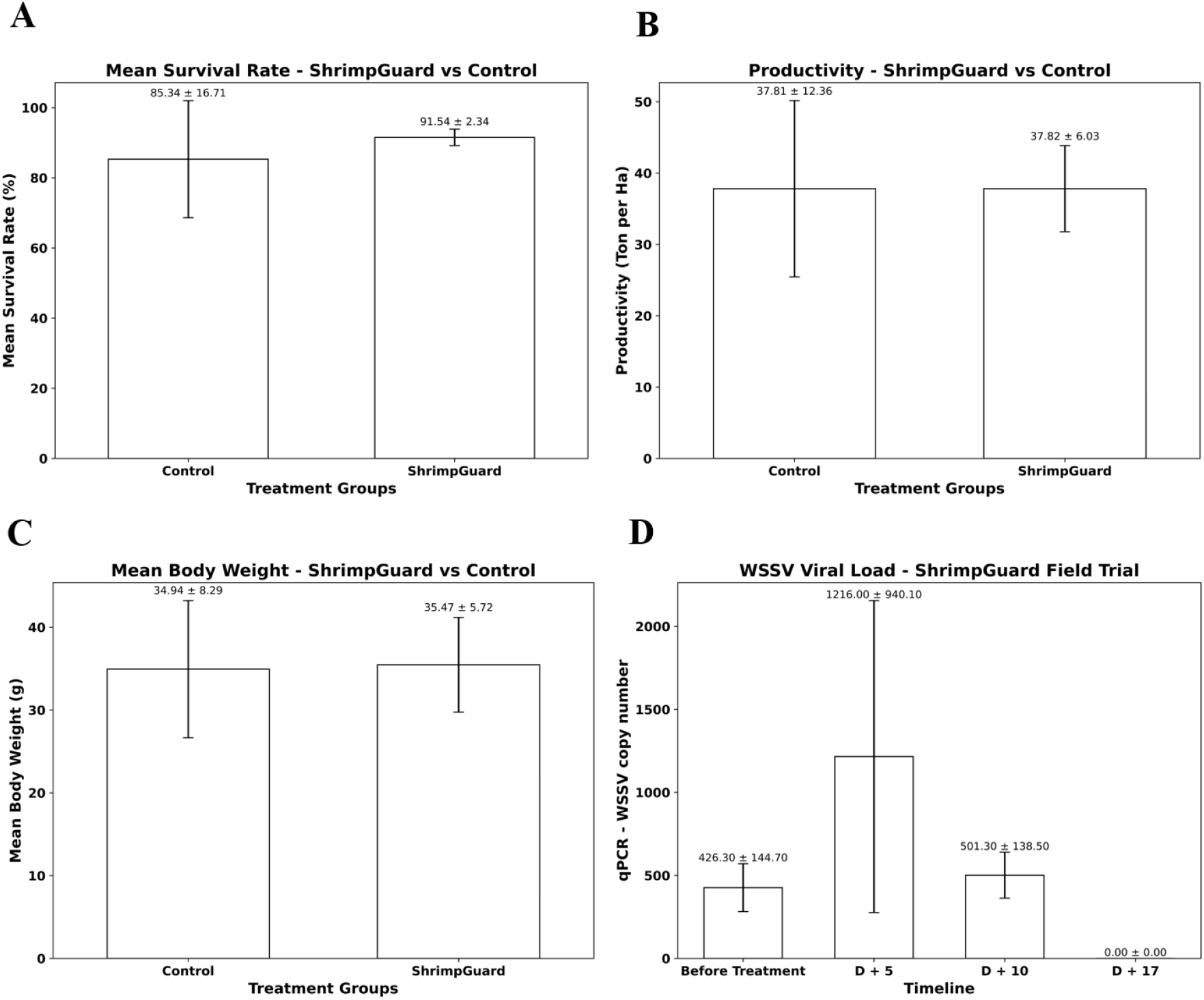
Efficacy evaluation of ShrimpGuardTM field trial: A) Mean survival rate, B) Productivity, and C) Mean body weight from the field trial-1 of ShrimpGuard treated and control treated shrimp grown in farm ponds. D) WSSV copy number at specific intervals of time from field trial-2. Survival rate, productivity, and average body weight are calculated at the day of harvest. Viral load was measured at specific intervals by qPCR in the field trial-2.

The survival rates of Group 3, 4 & 5 were 86.67%, 100% & 84.45 ± 3.14% respectively as shown in Fig. 5C. One-way ANOVA indicated a highly significant difference among the treatment groups (p < 0.0001). Tukey’s multiple comparison test revealed that all treatment groups—TW1, TW2, and TW3—showed significantly higher survival compared to the Positive Control group (p < 0.0001 for all comparisons) after 20 days post-challenge under lab conditions. Among the TW biologics, shrimps treated with TW-2 biologic exhibited the largest difference relative to the Positive Control - showed 100% survival under controlled WSSV challenge.

#### Oral trials: Efficacy trials with per os challenge

In this trial, each tank contained 6 shrimps per replicate. The shrimps were acclimatized in trial tanks for 1 day. After acclimatization, Group-1 (negative control, no virus, no TW biologic) & Group-2 (positive control, virus only no biologic) shrimps were fed with normal feed, while Groups 3, 4 & 5 were fed with feed top-coated with 2% TW biologics - TW-1, TW-2 & TW-3 respectively for 5 days. After the primary dose, all the shrimps were maintained under normal diet for 3 days. On the 10th & 11th day of the trial, all groups except Group-1 were infected with WSSV by per os challenge method (36–38, 41). After the WSSV challenge, all shrimps were monitored and maintained for 16 days under normal diet. On 17th day & 24th day of the trial the feed included oral administration of the respective treatment diets. The schematic of the trial plan is shown in Fig. 5B. The survival of shrimps were monitored for 16 days post challenge. After 16 days post-challenge, the survival rate of Group-1, which was not infected with virus and not fed with TW biologics, was 88.89 ± 9.62%. Group-2, which was infected with WSSV but not fed with TW biologics, experienced 100% mortality. The survival rates of Group 3, 4 & 5 were 72.22 ± 34.69%, 83.33 ± 16.67% & 11.11 ± 9.62% respectively as shown in Fig. 5D. One-way ANOVA showed a significant difference among the groups (p = 0.0008). Tukey’s multiple comparison test revealed that both TW-1 (p = 0.0072) and TW-2 (p = 0.0037*) groups differed significantly from the Positive Control, while TW-3 showed no significant difference (p = 0.778*) after 16 days post-challenge under lab conditions. Among the treatments, TW-2 demonstrated the largest mean difference from the Positive Control, indicating comparatively better performance - showed the highest survival rate with 83%. Until here, we have presented results of three TW biologics. However, to identify the best performing biologic, 8 additional TW biologics (TW-4 to TW-7 in Trial-S1A & TW-8 to TW-11 in Trial-S1B) were tested across two independent trials. The experimental design followed the same trial plan as the previous oral trial (schematic shown in Fig. 5B). In Trial S1A, after 10-days post WSSV challenge, the survival rates of treatment groups were: negative control group - 94.44 ± 9.62%, positive control group - 0%, TW-4 – 27.78 ± 34.69%, TW-5 – 22.22 ± 9.62%, TW-6 - 33.33 ± 44.1% & TW-7 – 33.33 ± 16.67%. In Trial S1B, after 10-days post challenge, the survival rates of treatment groups were: negative control group 94.44 ± 9.62%, positive control group – 11.11 ± 9.62%, TW-8 – 38.89 ± 19.25%, TW-9 – 38.89 ± 25.46%, TW-10 - 33.33% & TW-11 – 44.44 ± 9.62%. All TW-biologics from Trial S1A & S1B showed survival rate not significantly different from positive control (p > 0.1). Details of the trial design and corresponding data are provided in Supplementary information S1. These biologics delivered protection between 22.22% & 44.44%. As these TW biologics failed to meet our internally defined success criteria of 80% survival rate, they were not taken further.

Overall, results from the safety and efficacy trials demonstrated that the administration of TW-2 biologic provided a strong protective effect against WSSV. Following per os WSSV challenge & injection treatment, an 83% survival rate was observed over 10 days. This is only 5% less compared to tanks that were not treated with virus. When TW-2 was administered via top-coating with feed, it resulted in 100% survival for 20 days under a controlled intramuscular injection WSSV challenge, and 83% survival for 16 days under per os WSSV challenge. TW-2 has also demonstrated a protective effect in preliminary trials conducted in a different shrimp species, *Penaeus monodon* (Experimental procedures and trial data are provided in the Supplementary Information S2). Hence, we demonstrate that even in orally delivered biologics, there is specificity in the best performing solution, i.e. TW-2, consistent with the injection based trials. This is not caused by method of encapsulation, or any other factor as all other parameters other than the biologic remain consistent in the experiment group that are fed with TW biologics. Based on these results, TW-2, named ShrimpGuard™, was tested in field conditions.

## Discussion

WSSV is a major pathogen causing losses in the shrimp industry. Since its detection thirty years ago, multiple protein based and RNAi based solutions have been tested in shrimps as solutions for WSSV infections. RNAi technology incorporated in feed via encapsulation has been shown promising results in the laboratory (52). Various subunit protein and peptide-based solutions have been tested and demonstrated to be effective, particularly when administered via injection, as summarized comprehensively in the review by Feng et al. (2018) (30). Differences in production methods have been reported across various expression systems, including

E. coli, Bacillus subtilis, baculovirus, yeast and transgenic organisms. Among them E. coli is one of the most commonly used expression system and has produced effective WSSV vaccines, but it has been reported that bacterial expression systems are limited by the formation of insoluble inclusion bodies – leading to overall reduced efficacy (53). In addition, as most of the solutions are designed for injection-based delivery, it is not practical to implement these at large scale in commercial farms.

Recent studies indicate that insects have immune memory and specific immunity can be transmitted to their progenies. In a study in honeybees, researchers have shown that when parents are injected with immune-elicitors, their offspring exhibit improved protection (54). Another related study also demonstrated that this parental immunity is transferred via the egg yolk protein – vitellogenin (55, 56). Similar studies in Drosophila melanogaster have shown that immune priming through activation of the IMD pathway can produce immune memory and increase survival upon re-infection (57). Take together all these studies suggest that insects indeed have an immune system that can protect against future infection using immune memory. Similar to these systems, shrimps are also invertebrates, hence these protective mechanisms would exist in shrimps too. Results demonstrated here elucidated that indeed an immune response is observed in shrimps albeit it may not be through the conventional adaptive immune systems well studied in vertebrates.

Here we explore these possibilities and combine them with effective delivery systems to be able to bring industry ready solutions for major diseases. These studies done with ShrimpGuard™ collectively demonstrate the potential of ShrimpGuard™ to effectively manage WSSV infections in shrimp aquaculture. The laboratory trials demonstrated that the product is entirely safe to use and a survival rate of more than 80 % was observed upon WSSV infection in shrimps treated with it. On the contrary, under similar conditions of infection complete mortality is observed in shrimps not treated with ShrimpGuard™. ShrimpGuard™ was also evaluated for its performance in field conditions in a farm. Unlike laboratory trials, in field trials it is not possible to evaluate the efficacy of ShrimpGuard treated shrimp by actively subjecting them to WSSV challenge. Hence, evaluation of the in-field performance of anti-WSSV is based on the comparison of the multiple parameters, including % survival, weight gain, production per hectare, copy number of pathogens on random sampling, etc. to indicate that the product does not have any negative effect on shrimp yield. We compared the % survival, average weight, and production of the shrimp in the pond treated ShrimpGuard™ treated with a control pond which were not treated with it. Percentage survival post-harvest was 85.34 ± 16.71 % in control pond, and 91.54 ± 2.34 % in the ShrimpGuard™ treated pond. Average weight of shrimp in the control pond was 34.94 ± 8.29 gm and the ShrimpGuard™ treated was 35.47 ± 5.72 gm. Productivity of the control pond was estimated to be 37.81 ± 12.36 ton shrimp/hectare, and that of the Shrimp-Guard™ treated pond was estimated to be 37.82 ± 6.04 ton shrimp/hectare. These numbers unambiguously indicate that the ShrimpGuard™ can be used in the field without any negative effect on the farm yield. Upon use of ShrimpGuard™, there is a clear demonstration that the viral load in the ponds that were detected with WSSV did not increase. This is subject to early detection of the disease when the shrimp are still feeding and have not lost their appetite. Furthermore, given there is an observable trend towards improved survival rates, productivity, and average shrimp weight in ShrimpGuard™ treated ponds, this indicates that immunostimulant properties of ShrimpGuard™ have a positive impact on shrimp yield and growth.

Beyond ShrimpGuard™’s proven safety and efficacy, as an immunostimulant feed additive, it can be easily incorporated into existing feeding regimes along with other nutritional supplements. When used in combination with good aquaculture practices such as biosecurity, optimal water quality management, and the use of pathogen-free seed, Shrimp-Guard™ offers a sustainable strategy to improve shrimp health and resilience against WSSV infections.

## Acknowledgements

This work is supported by intramural funds from Teora Lifesciences Pvt Ltd. The authors wish to thank Prof. I.S. Bright Singh, National Centre for Aquatic Animal Health, Cochin University of Science Technology, for independently conducting some of the experimental trials associated with this study. The authors would also like to express their sincere gratitude to Dr. Krishna R. Salin, Associate Professor, and Dr. Ha Thanh Dong, Assistant Professor, in Aquaculture and Aquatic Resources Management, Department of Food, Agriculture, and Natural Resources, Asian Institute of Technology, Thailand and Dr. Megha K Bedekar, Principal Scientist & Head, Aquatic Environment & Health Management Division, ICAR – Central Institute of Fisheries Education – Mumbai, India for their valuable guidance on laboratory and field trials, as well as their overall support in matters related to aquatic health.

## Supplementary Information

### S1. Unsuccessful experimental challenge trial of TW biologics in P. vannamei against WSSV

The experimental design followed the same trial plan as the oral trial (schematic shown in Fig. 4B). Each trial comprised six treatment groups (each group in 3 replicates), including a negative control, a positive control, and four TW biologic treatments. The tables & graphs below show the survival data from preliminary unsuccessful lab trials conducted during the development phase of ShrimpGuard™. Each trial tested different candidate TW biologics. These studies don’t show significant improvement in survival, but were helpful in developing our end products.

**Figure S1A.**
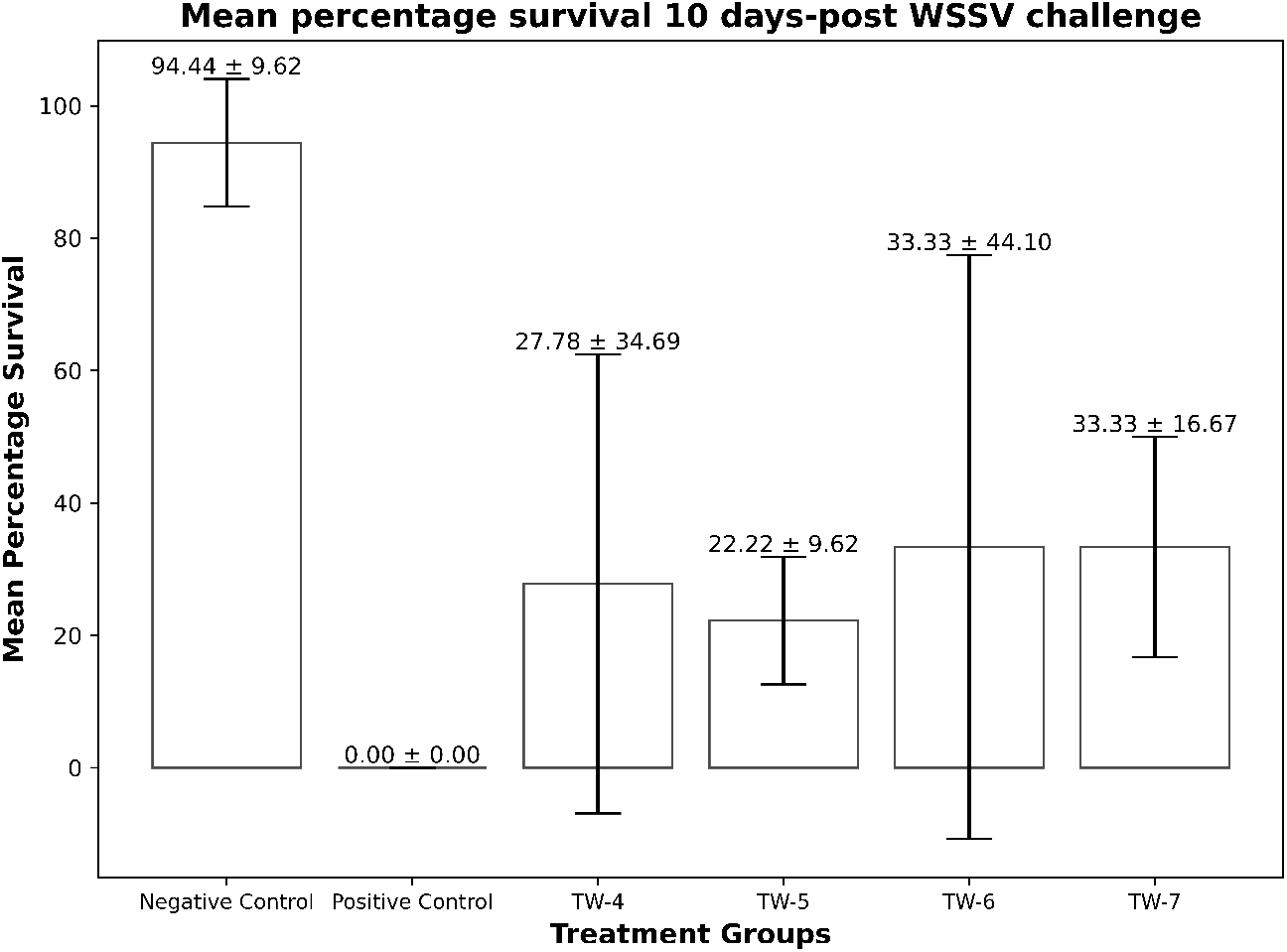
Efficacy assessment of TW biologics – TW-4, TW-5, TW-6 TW-7 against WSSV in P. vannamei. Percentage survival of shrimps 10 days post WSSV infection in the indicated treatment group. Negative control group was not infected with WSSV. The error bars represent the standard

#### Trial ID: S1A

In Trial S1A, after 10-days post challenge, the survival rates of treatment groups were: negative control group 94.44 ± 9.62%, positive control group - 0%, TW-4 – 27.78 ± 34.69%, TW-5 – 22.22 ± 9.62%, TW-6 - 33.33 ± 44.1% & TW-7 – 33.33 ± 16.67% respectively. One-way ANOVA indicated a significant overall difference among the groups (p = 0.0108). However, Tukey’s multiple comparison test revealed no significant pairwise differences between the Positive Control and any of the treatment groups (TW-4, TW-5, TW-6 or TW-7; all p > 0.5).

**Figure S1B.**
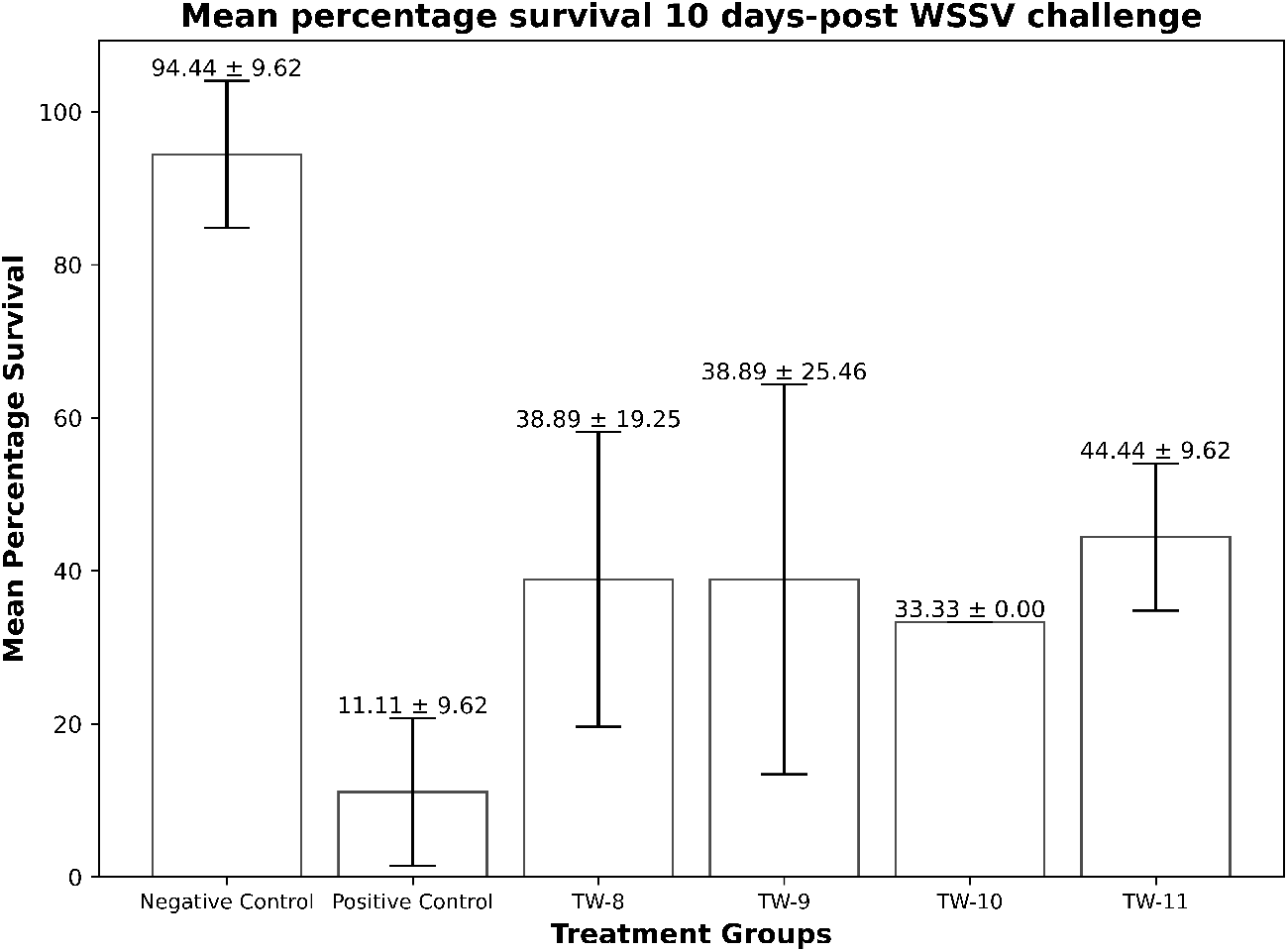
Efficacy assessment of TW biologics – TW-8, TW-9, TW-10 & TW-11 against WSSV in P. vannamei. Percentage survival of shrimps 10 days post WSSV infection in the indicated treatment group. Negative control group was not infected with WSSV. The error bars represent the standard deviation of % survival in three independent different tanks of a given shrimp group.

#### Trial ID: S1B

In Trial S1B, after 10-days post challenge, the survival rates of treatment groups were: negative control group 94.44 ± 9.62%, positive control group – 11.11 ± 9.62%, TW-8 – 38.89 ± 19.25%, TW-9 – 38.89 ± 25.46%, TW-10 - 33.33% & TW-11 – 44.44 ± 9.62% respectively. One-way ANOVA indicated a significant overall difference among the groups (p = 0.0005). However, Tukey’s multiple comparison test revealed no significant pairwise differences between the Positive Control and any of the treatment groups (TW-8, TW-9, TW-10 or TW-11; all p > 0.1).

### S2. Experimental challenge trial of TW biologics in *Penaeus monodon* against WSSV

To evaluate the potential protective efficacy of TW biologics against WSSV in *Penaeus monodon*, an experimental challenge trial was conducted under controlled laboratory conditions.

#### Trial Plan

*Penaeus monodon* PLs were divided into 5 treatment groups (each group comprising three tanks, with 10 PLs in each tank) – positive control, negative control, TW-1, TW-2 & TW-3. Prior to the challenge, PLs are fed their respective diets for 7 days. ShrimpGuard was incorporated into commercial shrimp feed at required inclusion rate for TW treatment groups other than control groups. On Day 7, all shrimp in TW treatment groups (TW-1, TW-2 & TW-3) and positive control group were challenged WSSV at a predetermined infectious dose previously tested to cause moderate to high mortality in *Penaeus monodon*. Shrimp were monitored for 12 days post-challenge. At the end of the trial, cumulative mortality & survival rate was compared between groups. TW-2 product was found to be promising, as the shrimps which survived the challenge were PCR negative to WSSV diagnostics.

#### Results

The TW-2 group showed a trend toward reduced cumulative mortality (36.67%) compared to the positive control group (60%). One-way ANOVA indicated a significant overall difference among the groups (p = 0.0058). However, Tukey’s multiple comparison test revealed no significant pairwise differences between the Positive Control and any of the treatment groups (TW-1, TW-2, or TW-3; all p > 0.05). While the protective effect of ShrimpGuard against WSSV, (63.3% survival rate) in *Penaeus monodon* did not reach statistical significance in this trial, observed trends in mortality reduction and delayed disease progression indicate a potential benefit.

**Figure S2.**
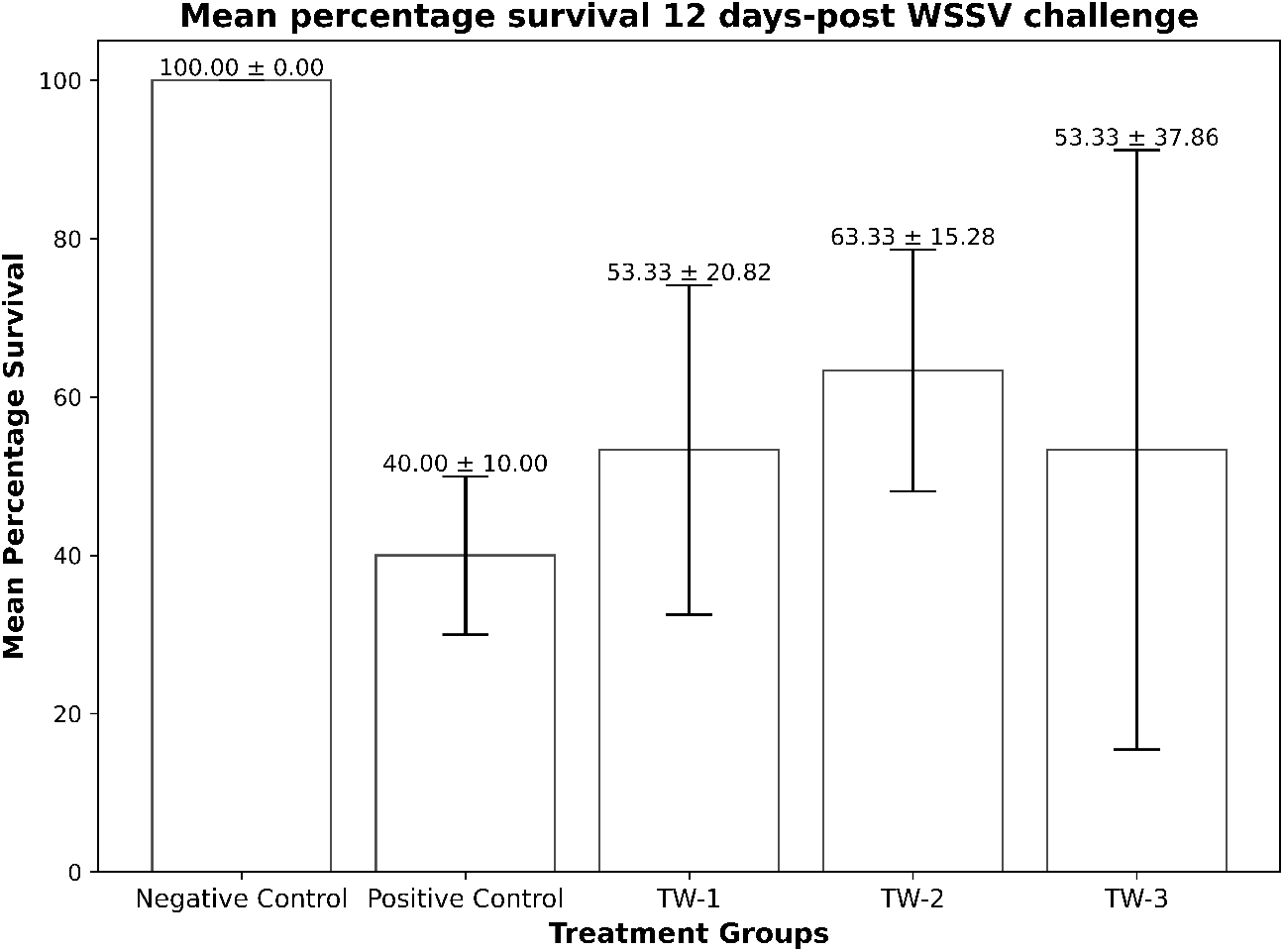
Efficacy assessment of TW biologics administered orally to *Penaeus monodon*. Percentage survival of shrimps 12 days post WSSV infection in the indicated treatment group. Negative control group was not infected with WSSV. The error bars represent the standard deviation of % survival in three independent different tanks of a given shrimp group

